# Amitosis confers benefits of sex in the absence of sex to *Tetrahymena*

**DOI:** 10.1101/794735

**Authors:** Hao Zhang, Joe A. West, Rebecca A. Zufall, Ricardo B. R. Azevedo

## Abstract

Sex appears to be the most successful reproductive strategy in eukaryotes despite its many costs. While a complete explanation for sex’s success remains elusive, several evolutionary benefits of sex have been identified. It is predicted that, by forgoing these benefits, asexual lineages are evolutionary dead-ends. Consistent with this prediction, many asexual lineages show signs of accelerated accumulation of deleterious mutations compared to their sexual relatives. Despite these low expectations, some asexual eukaryotic lineages appear to be successful, including the ciliate *Tetrahymena*. Here, we show that the mechanism of somatic nuclear division in *Tetrahymena*, known as amitosis, provides benefits similar to sex, allowing for the long-term success of asexual lineages. We found that, when compared to mitosis, amitosis with chromosome copy number control reduces mutation load deterministically, slows the accumulation of deleterious mutations under genetic drift, and accelerates adaptation. These benefits arise because, like sex, amitosis can generate substantial genetic variation in fitness among (asexual) progeny. Our results indicate that the ability of *Tetrahymena* to persist in the absence of sex may depend on non-sexual genetic mechanisms conferring benefits typically provided by sex, as has been found in other asexual lineages.

## Introduction

Although rare throughout ciliates, obligately asexual lineages are abundant, and possibly ancient, in the genus *Tetrahymena* (Doerder, 2014). The reason for this abundance is unknown. One possibility is that the peculiar genomic architecture of *Tetrahymena* allows it to avoid some of the negative consequences of asexuality (Doerder, 2014; Zufall, 2016).

Ciliates are microbial eukaryotes characterized by the separation of germline and somatic functions into two distinct types of nuclei within a single cell. The somatic macronucleus (MAC) is the site of all transcription during growth and asexual reproduction, and the germline micronucleus (MIC) is responsible for the transmission of genetic material during sexual conjugation (figure 1). Following conjugation, a zygotic nucleus divides and differentiates into the two types of nuclei (figures 1*A*, 1*B*). During this differentiation, the macronuclear genome undergoes massive rearrangements resulting in a genome with many small, highly polyploid, acentromeric chromosomes (Chalker, 2008). This genome structure results in amitotic macronuclear division (figures 1*C*, 1*D*).

**Figure 1:**
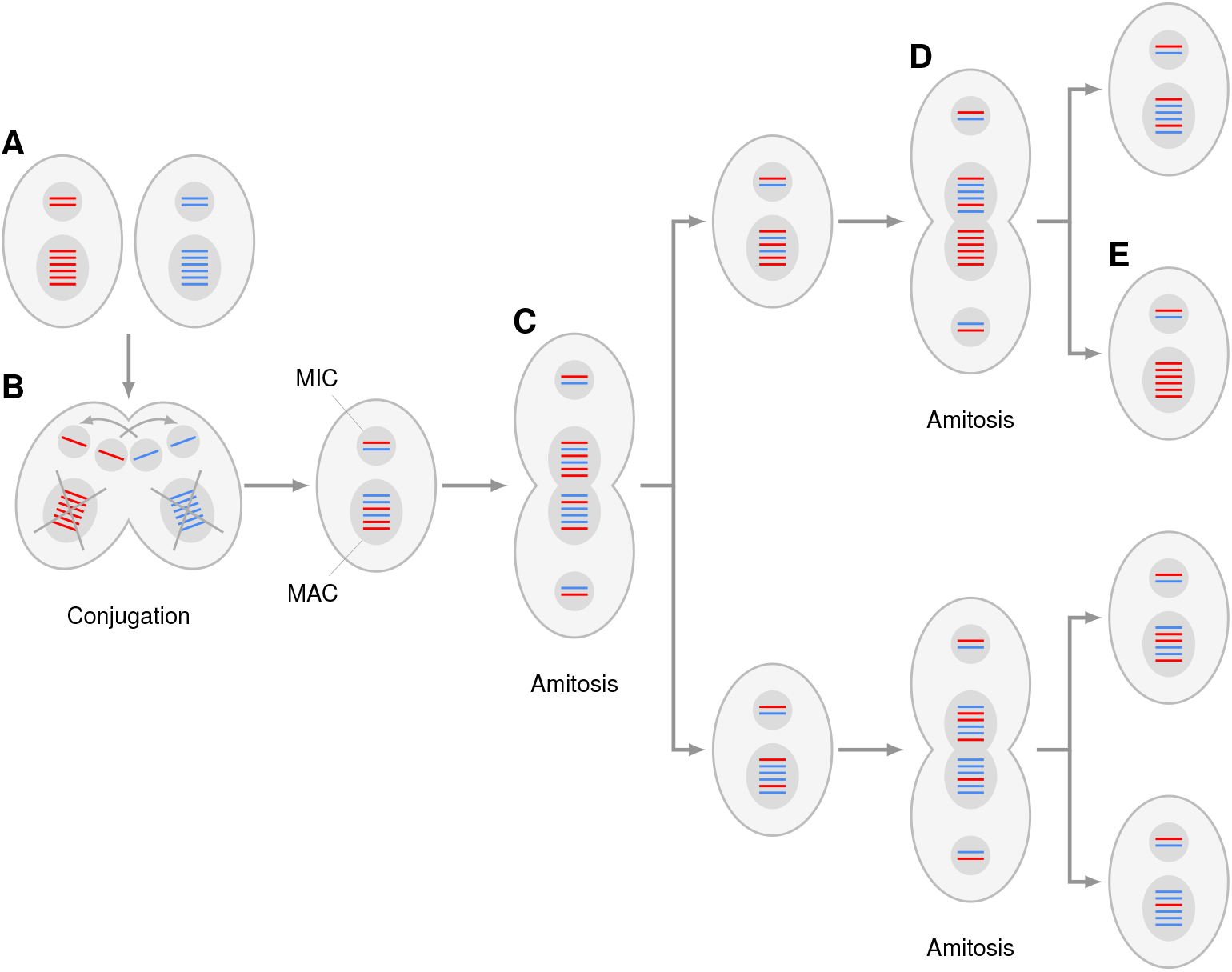
Amitosis with chromosome copy number control generates variation among individuals. Schematic of sexual conjugation followed by two rounds of asexual division. For simplicity, only one chromosome is shown: it occurs in two copies in the micronucleus (MIC) and six copies in the macronucleus (MAC) (in reality, each chromosome occurs in 45 copies in the *Tetrahymena thermophila* MAC). *A*, During sexual reproduction (conjugation), the diploid MIC undergoes meiosis (Jahn and Klobutcher, 2002; Orias et al., 2011). *B*, Two cells can fuse transiently and exchange haploid meiotic products. A resident meiotic product then fuses with the transferred meiotic product to produce a new diploid zygotic nucleus, which divides to generate the new MIC and MAC (the old MAC is destroyed). During asexual reproduction (*C, D*), the MIC divides by mitosis while the MAC divides by amitosis. Amitosis allows the random segregation of parental chromosomes among daughter cells generating variation among individuals. Ultimately, this results in phenotypic assortment, in which individual chromosomes in the MAC become completely homozygous within several generations (Doerder et al., 1992) (*E*). *T. thermophila*, has an unknown copy number control mechanism that results in an approximately equal number of homologous chromosomes in each daughter cell (Orias et al., 2011).

Amitosis generates variation among individuals in the number of copies of each allele at a locus. In most ciliates, amitosis results in differing numbers of chromosomes among progeny, which eventually leads to senescence and death (Bell, 1988). However, *Tetrahymena* have an unknown mechanism to control chromosome copy number during amitosis that results in roughly constant ploidy (Orias et al., 2011). A quarter of the 2,609 *Tetrahymena*-like wild isolates studied by Doerder (2014) lacked a MIC and were, therefore, asexual. To test whether amitosis with chromosome copy number control can account for the relative success of asexual *Tetrahymena*, we examined the evolutionary consequences of various forms of reproduction, nuclear division, and ploidy.

## Methods

### Model

#### Population

We model an infinite-sized population of asexual organisms of ploidy *n* reproducing in discrete generations. We begin by considering a single locus. The state of the population is given by a vector of frequencies 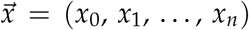, where *x*_*i*_ is the frequency of the genotype with *i* deleterious mutations and *n* − *i* wildtype alleles. Every generation, the population undergoes natural selection, mutation, and reproduction (mitosis or amitosis).

#### Natural selection

Natural selection causes the frequency of individuals with *i* deleterious mutations to change by 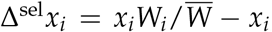, where *W*_*i*_ = 1 + *i s*_*d*_/*n* is the fitness of an individual with *i* deleterious mutations, *s*_*d*_ *<* 0 is the effect of a deleterious mutation in a homozygous state, and 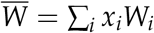 is the mean fitness.

#### Mutation

An individual with *i < n* deleterious mutations will mutate into an individual with *i* + 1 mutations with probability *µ*_*d*_(*n* − *i*) where *µ*_*d*_ is the deleterious mutation rate per wildtype allele per generation. We assume that all mutations are deleterious and irreversible. Thus, mutation will change *x*_*i*_ by

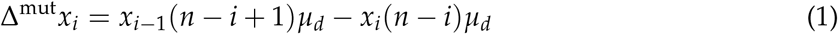

#### Reproduction

Mitosis has no effect on genotype frequencies: Δ^mit^ *x*_*i*_ = 0. An individual with *j* deleterious mutations reproducing by amitosis has offspring with *i* mutations with probability (Schensted, 1958)

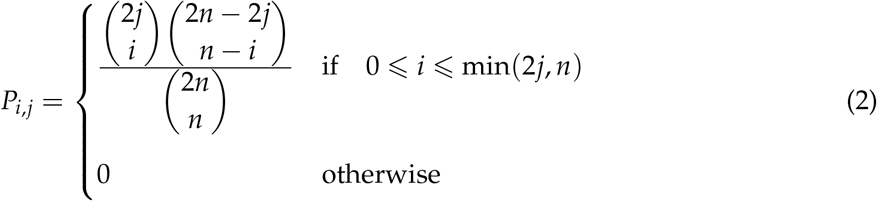

Thus, amitosis changes *x*_*i*_ by 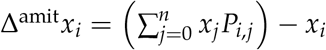.s

#### Evolution

We assume that these population genetic processes operate independently every generation. Thus, evolution under reproductive strategy *ρ* (e.g., amitosis) is described by

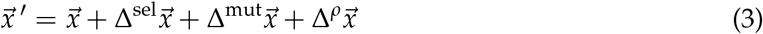

where 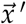 are the genotype frequencies in the next generation. See equation (A1) for an example.

#### Equlibrium

A population of individuals with *x*_0_ *>* 0 evolving according to equation (3) evolves towards a stable equilibrium where the mean fitness is *Ŵ* _*ρ*,1_. If there are *L* fitness loci with the same *µ*_*d*_ and *s*_*d*_, and there is linkage equilibrium between these loci, the mean fitness at equilibrium will be *Ŵ*_*ρ,L*_ =(*Ŵ*_*ρ*,1_)^*L*^. Appendix A shows the derivation of *Ŵ* _*ρ,L*_ for amitosis, *Ŵ*_amit_, in diploids (*n* = 2).

When ploidy was *n >* 2, *Ŵ*_amit_ was calculated numerically by iterating equation (3). Equilibrium was inferred when the Euclidean distance between consecutive *x* was smaller than 10^−6^. An approach similar to that used by Kondrashov (1994) to model vegetative reproduction should allow the derivation of general analytical results for *n >* 2.

### Stochastic simulations

Stochastic, individual-based simulations were conducted within a Wright-Fisher framework (Ewens, 2004). Individuals have *L* fitness loci and undergo a mutation–selection–reproduction life cycle. Populations have constant size *N*. Initially, all individuals are mutation-free and have a fitness of *W* = 1.

Every generation, each individual may acquire a new mutation at a fitness locus with the probabilities shown in equation (1). An individual can acquire multiple mutations, but only one per locus.

Under asexual reproduction (mitosis or amitosis), *N* individuals are chosen at random to reproduce, with replacement, with probability proportional to their fitness. Each individual chosen to reproduce is allowed to generate one offspring. Under mitosis, the offspring is an exact copy of the parent; under amitosis, the number of mutant alleles inherited by the offspring at each locus is drawn at random with probability given by equation (2). The parents are discarded after reproduction.

Under sexual reproduction, 2*N* individuals are chosen randomly with replacement, with probability proportional to their fitness. We then create *N* pairs of individuals from this set at random without replacement. Each pair is allowed to generate one offspring with free recombination among the *L* loci. The parents are discarded after reproduction.

The rate of accumulation of drift load was measured as the slope of a linear regression of population ln 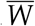 against generation. Equilibrium was evaluated using the slope of a linear regression of population 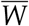 against generation. Equilibrium was inferred when the average slope was not statistically significantly different from zero over a large number of replicate populations. The slopes were evaluated between generations 300 and 600 in diploids and between generations 9 × 10^3^ and 10^4^ in 45-ploids.

### Code availability

Mathematical analyses and numerical calculations were done using Mathematica 12.2. Stochastic simulations were performed using software written in Python 3.7. All code and data have been deposited in the in the Dryad Digital Repository (https://datadryad.org/stash/share/EyyVxUGkVJidWpo4wkMiWMBAEsFZtxg8Z3vtzK3Wz8s).

## Results and Discussion

Most mutations with effects on fitness are deleterious but natural selection cannot remove all of them from populations. As a result, many individuals carry deleterious mutations that reduce their fitness, which leads to a reduction in the mean fitness of populations, or mutation load. We begin by investigating the extent to which amitosis with chromosome copy number control affects mutation load.

Kimura and Maruyama (1966) showed that a population of asexual unicellular organisms reproducing by mitosis is expected to show the following mean fitness at equilibrium:

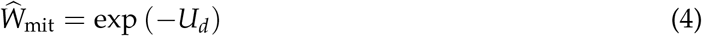

where *U*_*d*_ is the deleterious mutation rate per genome per generation. Equation (4) assumes that all mutations are (i) deleterious and (ii) irreversible, and that (iii) the population is very large, so we can ignore genetic drift. Equation (4) is valid for any ploidy.

In contrast, if an asexual diploid population reproduces by amitosis, its mean fitness at equilibrium is given by

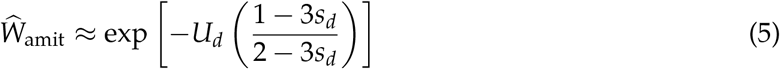

where *U*_*d*_ = 2*Lµ*_*d*_, *L* is the number of loci influencing fitness, *µ*_*d*_ is the deleterious mutation rate per locus per generation, and *s*_*d*_ *<* 0 is the effect on fitness of a deleterious mutation in a homozygous state (see Appendix A for derivation). Equation (5) relies on six additional assumptions: *µ*_*d*_ is (i) low and (ii) equal across loci; (iii) there is linkage equilibrium among fitness loci; (iv) all mutations have the same deleterious effect *s*_*d*_, and contribute to fitness (v) additively within loci (i.e., are codominant) and (vi) multiplicatively among loci (i.e., do not interact epistatically). The scenario described by equation (5) is purely theoretical because no diploid nucleus is known to reproduce amitotically. However, it allows us to compare the evolutionary consequences of amitosis to those of mitosis.

Equations (4) and (5) show that amitosis can reduce mutation load compared to mitosis in diploid populations. For example, if *U*_*d*_ = 0.1 and *s*_*d*_ = −0.1, the mean fitness at equilibrium is *Ŵ*_mit_ = 0.905 under mitosis and *Ŵ*_amit_ = 0.945 under amitosis. Thus, amitosis has a selective advantage over mitosis of *Ŵ*_amit_/*Ŵ*_mit_ − 1 = 4.4%. The deleterious mutation rate, *U*_*d*_, has a large effect on the benefit of amitosis: doubling the value of *U*_*d*_ more than doubles the advantage of amitosis to 9.1% (figure 2*A*). The selection coefficient of a deleterious mutation, *s*_*d*_, however, has a comparatively small effect on the benefit of amitosis: making mutations one tenth as deleterious (*s*_*d*_ = −0.01) causes the advantage of amitosis to increase to only 5.0% (figure 2*B*).

**Figure 2:**
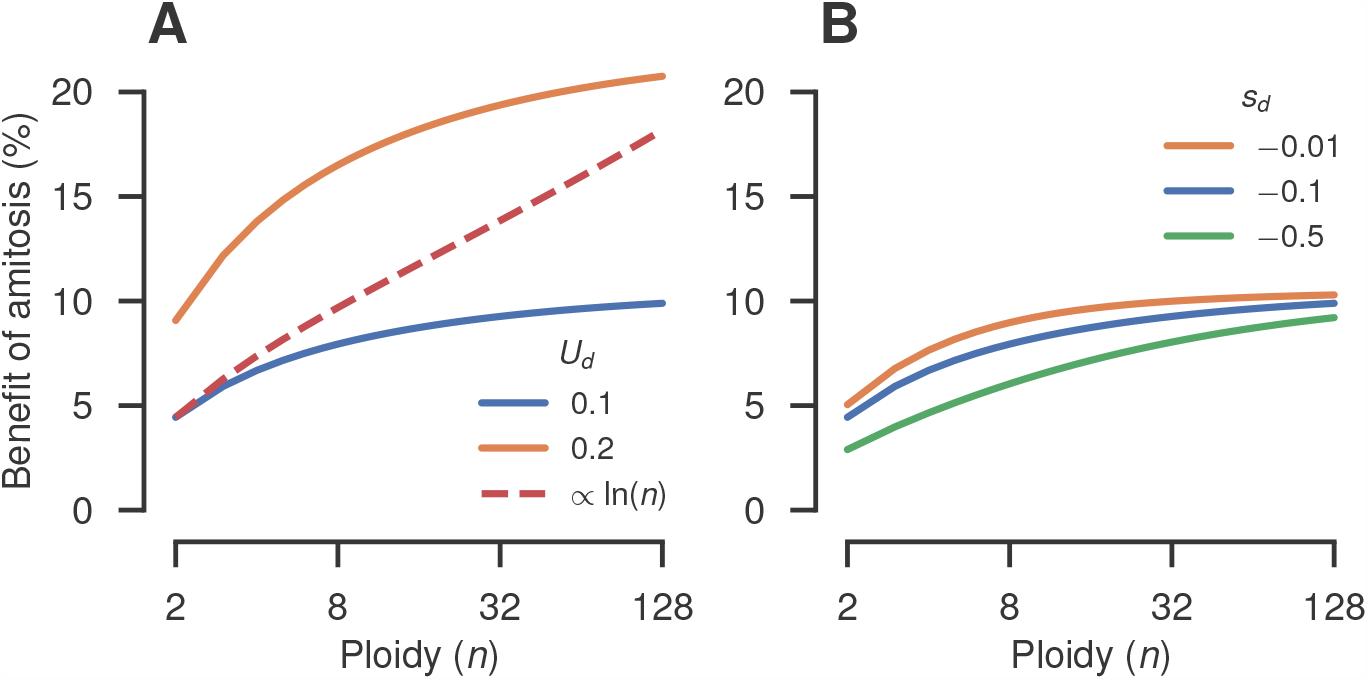
Amitosis with chromosome copy number control reduces mutation load relative to mitosis in large populations. Values show the selective advantage of amitosis over mitosis, *Ŵ*_amit_/*Ŵ*_mit_ − 1, at different ploidies (*Ŵ*_*ρ*_ is the mean fitness at equilibrium of a population of individuals following reproductive strategy *ρ* for a certain ploidy). *A*, Effect of the genomic deleterious mutation rate, *U*_*d*_. Solid lines show selective benefits corresponding to constant values of *U*_*d*_ at all ploidies. The dashed line assumes that a doubling of the ploidy results in a 10% increase in *U*_*d*_. Mutations have a deleterious effect of *s*_*d*_ = −0.1 at all ploidies. *B*, Effect of the selection coefficient of a deleterious mutation, *s*_*d*_. We set *U*_*d*_ = 0.1 at all ploidies. In both *A* and *B* we assumed that there were *L* = 100 fitness loci. Note that ploidy is shown in a log scale.

Amitosis with copy number control is observed in the genus *Tetrahymena*, which have high ploidy in their macronuclear genome (e.g., *T. thermophila* are 45-ploid). Interestingly, the benefit of amitosis relative to a mitotically reproducing organism with the same ploidy increases with ploidy (figure 2). For example, if *U*_*d*_ = 0.1 and *s*_*d*_ = −0.1, the benefit of amitosis increases to 6.7% in tetraploids, 7.9% in octoploids, 8.7% in 16-ploids, and so on. Further increases in ploidy cause diminishing returns in the benefit of amitosis. These expected benefits are conservative because they assume that the deleterious mutation rate, *U*_*d*_, is constant across ploidies. If, for example, doubling ploidy causes an increase of 10% in *U*_*d*_, a substantially greater benefit of amitosis would be achieved at high ploidies (figure 2*A*, dashed line). A mutation accumulation study estimated that *T. thermophila* has a deleterious mutation rate in the MIC of 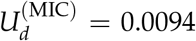 per genome per generation and that mutations have an expected deleterious effect of 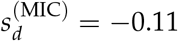 in a homozygous state (Long et al., 2016). If we assume that the MAC genome has 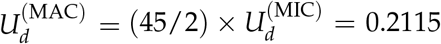 and 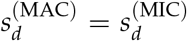, we estimate that amitosis has a benefit of 21.0% relative to mitosis in this species.

The analyses so far have ignored the effect of genetic drift. Drift can cause a population to accumulate deleterious mutations stochastically, further increasing genetic load, or drift load (Kimura et al., 1963; Crow, 1970; Poon and Otto, 2000). In asexuals this phenomenon is known as Muller’s ratchet (Muller, 1964; Felsenstein, 1974; Haigh, 1978). We now evaluate the extent to which amitosis with copy number control can slow down the accumulation of drift load. Populations of *N* = 10 or 100 diploid mitotic individuals experience strong Muller’s ratchet when *U*_*d*_ = 0.1 and *s*_*d*_ = −0.1 (figure 3*A*). Increasing population size to *N* = 10^3^ individuals causes the ratchet to slow down considerably, allowing populations to achieve mutation-selection equilibrium (figure 3*A*). Reproduction through amitosis makes populations less susceptible to Muller’s ratchet. The accumulation of drift load slows down by 39% (95% confidence interval, CI: 31%, 46%) in diploid populations of *N* = 10 individuals, and effectively halts in populations of *N* = 100 individuals (figure 3*C*).

**Figure 3:**
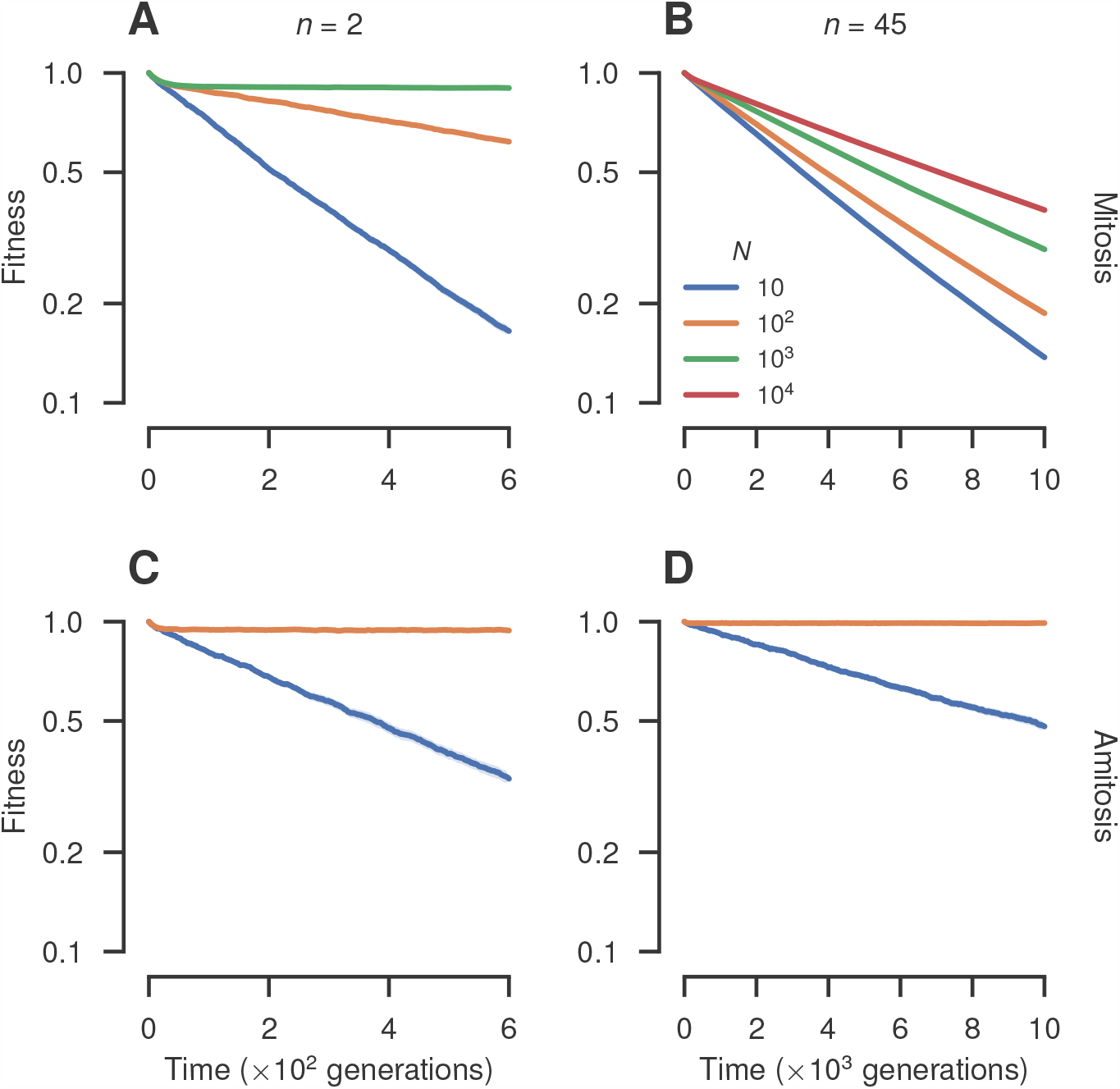
Amitosis with chromosome copy number control slows down the accumulation of drift load relative to mitosis. Evolutionary responses of mean fitness in populations of different sizes (*N*) and plodies (*n*), following different reproductive strategies. Lines show the means of stochastic simulations of 100 populations; shaded regions represent 95% CIs. *A*, Mitosis in diploids (*n* = 2). *B*, Mitosis with a ploidy of *n* = 45. *C*, Amitosis in diploids (*n* = 2). *D*, Amitosis with a ploidy of *n* = 45. We assumed *L* = 100 fitness loci, a genomic deleterious mutation rate of *U*_*d*_ = 0.1 per generation, that mutations have a deleterious effect of *s*_*d*_ = −0.1 in a homozygous state, and that, initially, all individuals are unmutated. Note that fitness is shown in a log scale.

The benefit of amitosis in slowing down the accumulation of drift load, like the deterministic benefit, increases with ploidy. Muller’s ratchet operates in populations as large as *N* = 10^4^ mitotic 45-ploid individuals (figure 3*B*). Amitosis is able to halt the accumulation of drift load in populations with as few as *N* = 100 45-ploid individuals (figure 3*D*). Even when amitotic populations are small enough to accumulate drift load, they do so more slowly than mitotic ones. For example, populations of *N* = 10 amitotic 45-ploid individuals accumulate drift load 64% (95% CI: 59%, 68%) more slowly than mitotic populations of the same size (figures 3*B*, 3*D*).

The benefits of amitosis over mitosis identified so far are analogous to benefits of sexual over asexual reproduction. In diploids, sexual reproduction by selfing confers a deterministic advantage over mitosis almost identical to that of asexual amitosis shown in equation (5) (Charlesworth et al., 1990*a*,*b*).

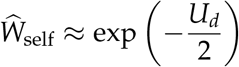

Unlike amitosis, sex with random mating in diploids only confers a deterministic advantage over asexual reproduction if there is negative epistasis between deleterious mutations (Kondrashov, 1988; Otto and Feldman, 1997), or if deleterious mutations are partially recessive (Chasnov, 2000; Otto, 2003). Sex can also counteract Muller’s ratchet (Muller, 1964; Felsenstein, 1974), much like amitosis (figures 3*A*, 3*C*). Are the benefits of asexual amitosis also similar to those of sexual reproduction when ploidy is high? We investigated this question in populations of *N* = 20 individuals of a 45-ploid organism like *T. thermophila* experiencing *U*_*d*_ = 0.1 and *s*_*d*_ = −0.1. Amitosis slows down the accumulation of drift load relative to mitosis by 90% (95% CI: 88%, 92%; figure 4*A*).

**Figure 4:**
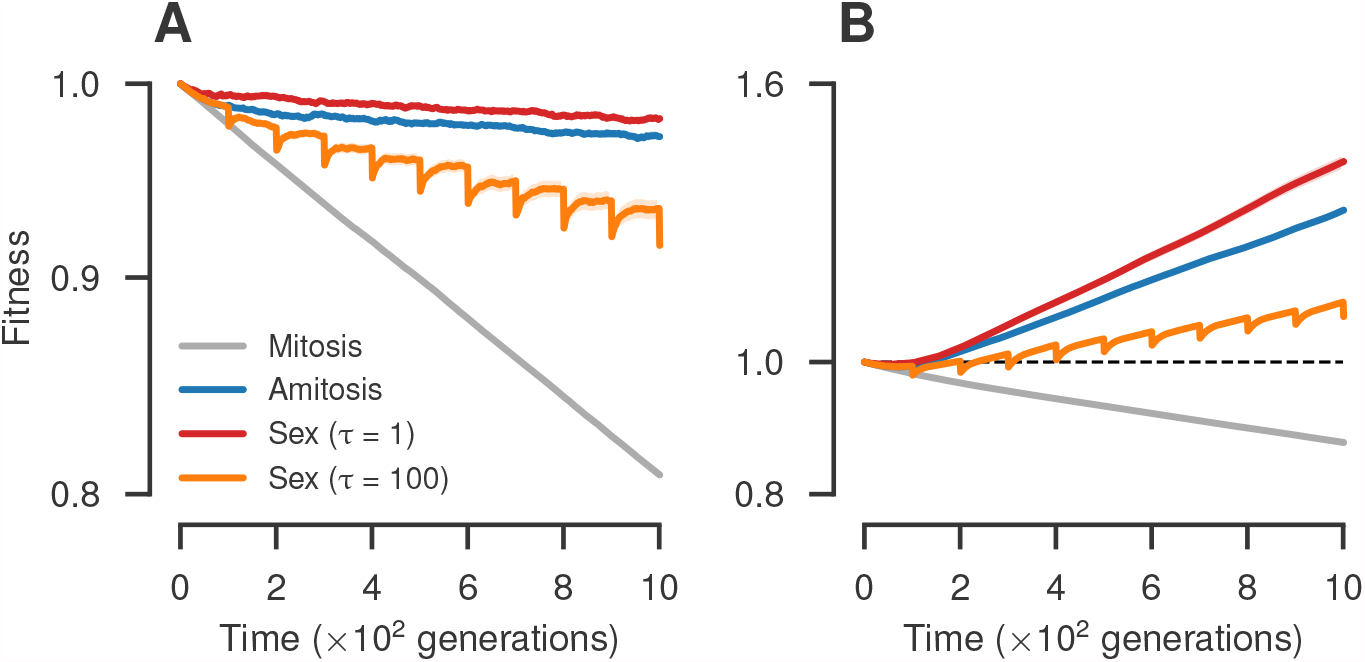
The benefit of amitosis with chromosome copy number control is similar to that of sex. Evolutionary responses of population mean fitness under different reproductive strategies. Lines show the means of stochastic simulations of 500 populations; shaded regions represent 95% CIs. *A*, Populations of *N* = 20 individuals with a deleterious mutation rate of *U*_*d*_ = 0.1 per genome per generation. All mutations are deleterious and have a selection coefficient of *s*_*d*_ = −0.1 in a homozygous state. *B*, Populations of *N* = 103 individuals with a genomic mutation rate of *U* = 0.1 per generation; 99% of mutations are deleterious and 1% are beneficial with selection coefficients of *s*_*d*_ = −0.1 and *s*_*b*_ = 0.1, respectively. We assumed that individuals have a MAC ploidy of *n* = 45 with *L* = 100 fitness loci, and that, initially, they carry no mutations. Sexual reproduction takes place with random mating and free recombination every *τ* generations. Note that fitness is shown in a log scale.

An organism like *T. thermophila* but reproducing sexually, with outcrossing, every generation (i.e., obligate sex with no amitosis) and then generating a 45-ploid macronucleus from the recombinant diploid micronucleus (see figure 1*A, B*) would slow down the accumulation of drift load by 92% (95% CI: 90%, 94%; *τ* = 1, figure 4*A*). However, *T. thermophila* cannot reproduce sexually every generation; rather, it requires approximately 100 asexual cell divisions to reach sexual maturity (Doerder et al., 1995; Nanney et al., 1955). Facultative sex every *τ* = 100 generations slows down the ratchet by only 68% (95% CI: 64%, 72%; measured based on fitness in the generation immediately before the population reproduces sexually), much less than amitosis (figure 4*A*). The benefit of amitosis is also comparable to that of sex in larger populations in the presence of beneficial mutations. In an evolutionary scenario under which asexual populations are not able to adapt, both amitosis and obligate sex every generation (*τ* = 1) allow populations to adapt, and more rapidly than facultative sex every *τ* = 100 generations (Figure 4*B*).

The results shown in figure 4 raise the intriguing possibility that amitosis is actually evolutionarily superior to facultative sex in *T. thermophila* and its relatives, which have *τ* ≈ 100. If true, this would lead to the prediction that asexual lineages should outcompete sexual ones in *Tetrahymena*. This could explain why obligately asexual lineages are abundant in *Tetrahymena* (Doerder, 2014). If this explanation is correct, we would expect that asexual lineages of *Tetrahymena* do not show the typical signs of accelerated accumulation of deleterious mutations compared to their sexual relatives (Paland and Lynch, 2006; Johnson and Howard, 2007; Neiman et al., 2010; Henry et al., 2012; Tucker et al., 2013; Hollister et al., 2015).

The hypothesis outlined in the previous paragraph may be invalid for two reasons. First, our analysis may overestimate the benefit of amitosis relative to facultative sex. Our hypothesis assumes that chromosome copy number control during amitosis is perfect, or at least, highly precise on an evolutionary time-scale. However, the precision of copy number control is unknown even in *T. thermophila*. Control of chromosome copy number could be less precise than we have assumed and, therefore, confer a smaller benefit to *Tetrahymena*. Second, our analysis may underestimate the benefit of facultative sex relative to amitosis. We have considered only two possible benefits of sex, both “mutational” in nature (Kondrashov, 1993). Other benefits of sex are not guaranteed to show the same pattern. For example, we have not considered the potential benefits of sex in the face of biotic interactions (Bell, 1982; Hamilton et al., 1990; Otto and Nuismer, 2004). Even if our hypothesis is correct, it is also conceivable that there are additional factors contributing to the relative success of asexual *Tetrahymena*. For example, it has been proposed that high ploidy alone may inhibit the accumulation of deleterious mutations through gene conversion (Maciver, 2016). However, this proposed advantage has not been modelled, and therefore it is difficult to evaluate.

What is the mechanistic basis of the benefits of amitosis identified here? The main difference between the two types of nuclear division is that amitosis, like sex, can generate more genetic variation in fitness than mitosis. For example, an *n*-ploid individual (we assume *n* is even for simplicity) with *n*/2 wild-type alleles and *n*/2 deleterious alleles will have a fitness of *W* = 1 − *s*_*d*_/2. Mutation will generate a variance in fitness of

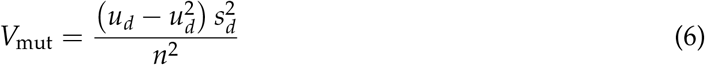

every generation, where *u*_*d*_ = *nµ*_*d*_/2 is the deleterious mutation rate at the locus per generation. Mitosis is not expected to generate any variance in fitness in addition to mutation (i.e., *V*_mit_ = *V*_mut_). Amitosis will, however, increase the variance in fitness further

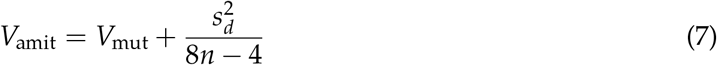

every generation (Schensted, 1958). Since *u*_*d*_ is likely to be low, amitosis is expected to increase the variance in fitness to a much greater extent than mutation, and therefore mitosis (*V*_amit_ » *V*_mit_). We propose that amitosis causes an increase in the additive genetic variance in fitness, therefore making natural selection more efficient—an analog of Weismann’s hypothesis for the advantage of sex (Weismann, 1887; Kondrashov, 1993; Burt, 2000). Consistent with this idea, the variance in fitness generated by amitosis relative to mitosis increases approximately linearly with ploidy according to equations (6) and (7)

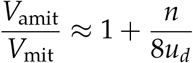

which explains why the benefit of amitosis relative to mitosis increases with ploidy.

We conclude that amitosis with chromosome copy number control confers benefits of sex in the absence of sex and can account for the high incidence of obligately asexual lineages in *Tetrahymena* (Doerder, 2014). Other successful asexual lineages also appear to show benefits of sex in the absence of sex albeit through different mechanisms (Gladyshev et al., 2008; Flot et al., 2013; Seidl and Thomma, 2014; Maciver, 2016).

## Acknowledgments

We thank P. Doerder, M. Orive, T. Paixão, and A. Kondrashov for comments on an earlier version of the manuscript, and E. Kelleher for discussions. R.A.Z. and R.B.R.A. acknowledge support from grant R01GM101352 from the National Institutes of Health. R.A.Z. acknowledges support from grant DEB-1911449 from the National Science Foundation (NSF). R.B.R.A. acknowledges support from grants DEB-1354952 and DEB-2014566 from the NSF. We acknowledge the use of the Maxwell and Opuntia clusters and the advanced support from the Research Computing Data Core at the University of Houston.

## Appendix A: Amitosis in diploids

### Model

A population of diploid individuals reproducing by amitosis evolves at one fitness locus according to the system of recursion equations defined by equation (3)

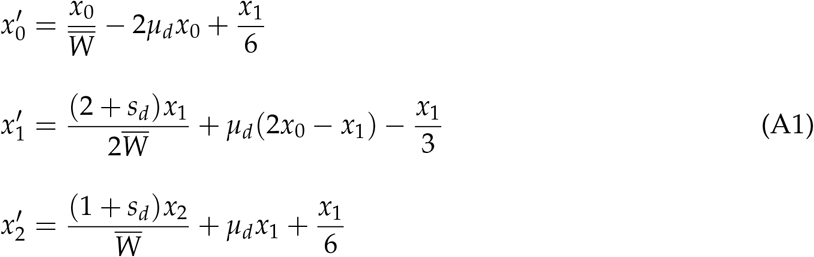

where the mean fitness of the population is

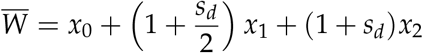

*x*_*i*_ is the frequency of the genotype with *i* deleterious mutations and *n* − *i* wildtype alleles, *µ*_*d*_ is the deleterious mutation rate per wildtype allele per generation, and *s*_*d*_ *<* 0 is the effect of a deleterious mutation in a homozygous state. Note that ∑_*i*_ *x*_*i*_ = 1.

#### Equilibrium

There is one equilibrium where unmutated individuals are present in the population (*x*_0_ *>* 0)

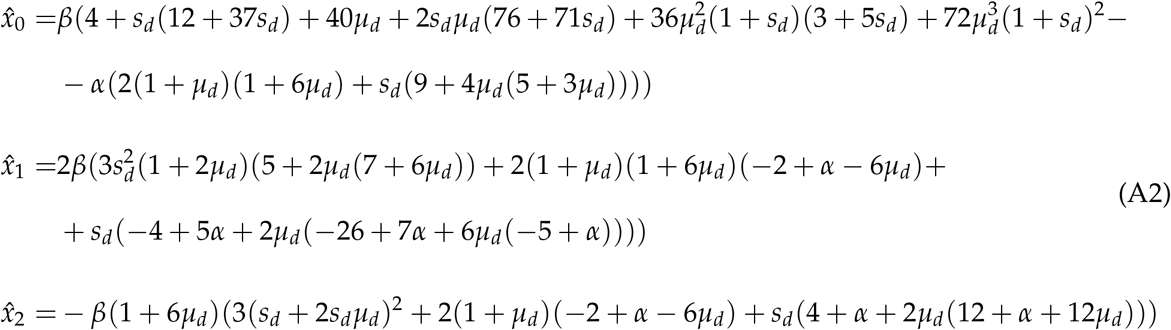

where

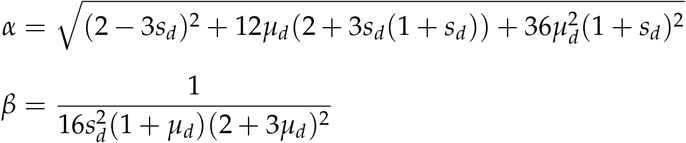

#### Stability of the equilibrium

The Jacobian matrix of the system is

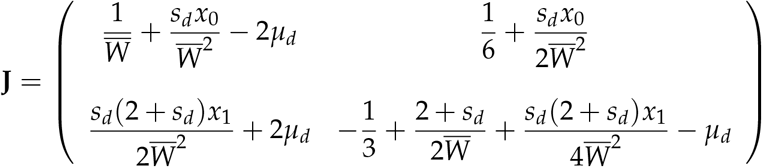

(We only need to consider *x*_0_ and *x*_1_ because *x*_2_ = 1 − *x*_0_ − *x*_1_.)

The characteristic equation of **J** evaluated at the equilibrium in equation (A2) is

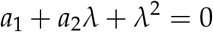

with coefficients

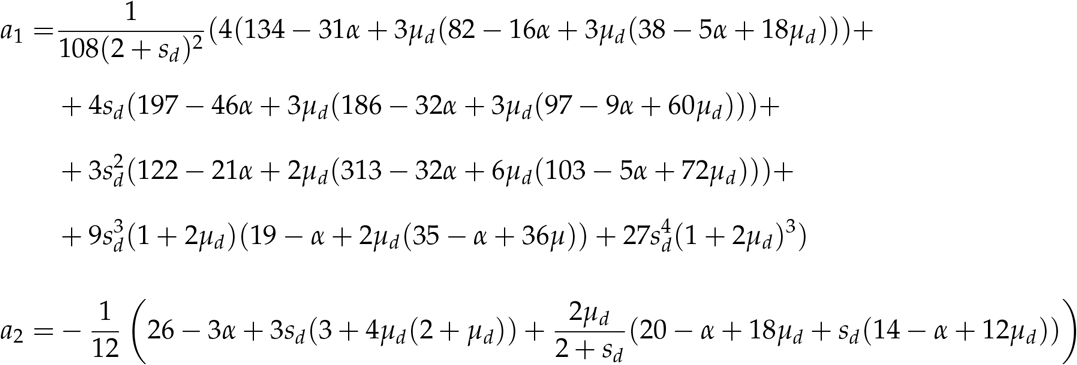

The equilibrium in equation (A2) is stable if the following Routh-Hurwitz conditions are met (Otto and Day, 2007)

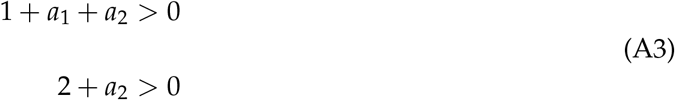

If we assume that mutations occur at a low rate (0 *< µ*_*d*_ ≪ 1) and are deleterious (−1 ≤ *s*_*d*_ *<* 0), the conditions in equation (A3) are met when

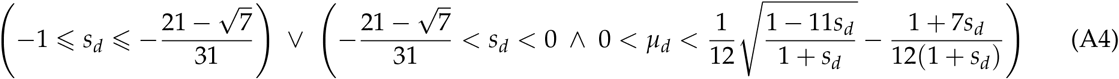

The equilibrium in equation (A2) is valid (∀_*i*_ : 0 ⩽*x*_*i*_ ⩽ 1) when the condition in equation (A4) is met.

#### Mean fitness at equilibrium

The mean fitness at the equilibrium defined by equation (A2) is

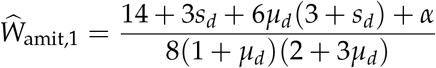

If there are *L* fitness loci with the same *µ*_*d*_ and *s*_*d*_, and there is linkage equilibrium between these loci, the mean fitness at equilibrium will be *Ŵ* _amit,*L*_ = (*Ŵ* _amit,1_)^*L*^. Taking a first-order Taylor expansion of ln (*Ŵ*_amit,*L*_) around *µ*_*d*_ = 0 we get equation (5).

An approach similar to that used by Kondrashov (1994) to model vegetative reproduction should allow the derivation of general analytical results for *n >* 2.

